# miCloud: a plug and play, on-premises bioinformatics cloud, providing seamless integration with Illumina genome sequencers

**DOI:** 10.1101/209734

**Authors:** Baekdoo Kim, Thahmina Ali, Konstantinos Krampis, Changsu Dong, Bobby Laungani, Claudia Wultsch, Carlos Lijeron

**Affiliations:** Weill Cornell Medicine - Hunter College Belfer Research Building, New York, NY; Department of Biological Sciences, Hunter College of The City University of New York, NY; American Museum of Natural History, New York, NY.

## Abstract

Benchtop genome sequencers such as the Illumina MiSeq or MiniSeq [1], [2] are revolutionizing genomics research for smaller, independent laboratories, by enabling access to low-cost Next Generation Sequencing (NGS) technology in-house. These benchtop genome sequencing instruments require only standard laboratory equipment, in addition to minimal time for sample preparation. However, post-sequencing bioinformatics data analysis still presents a significant bottleneck, for research laboratories lacking specialized software and technical data analysis skills on their teams. While bioinformatics computes clouds providing solutions following a Software as a Service (SaaS) are available ([3]–[6], review in [7]), currently, there are only a few options which are user-friendly for non-experts while at the same time are also low-cost or free. One primary example is Illumina BaseSpace [8] that is very easy to access by non-experts, and also offers an integrated solution where data are streamed directly from the MiSeq sequencing instrument to the cloud. Once the data is on the BaseSpace cloud, users can access a range of bioinformatics applications with pre-installed algorithms through an intuitive web interface. Nonetheless, BaseSpace can be a costly solution as a yearly subscription depending on whether the user is associated with an academic or private institution, ranges in price from $999 - $4,999. Additional “iCredits” [9] might need to be purchased for frequent users that exhaust the base credit allowance as part of the subscription. Considering the reduction of computer hardware cost in recent years, a multi-core Intel Xeon server with 64 GigaByte (GB) of memory and multiple TeraByte (TB) of storage is priced less than the yearly subscription to Basespace [10], and similarly when compared to renting compute cycles from providers such as Amazon Web Services (AWS) [11]. Furthermore, the current generation of laptops usually come with 6–10 GigaBytes (GB) of memory and 1 TeraByte (TB) of storage, providing enough computational capacity to analyze data from small NGS experiments [12] that include only a few samples.

---

We developed miCloud, a bioinformatics platform for NGS data analysis, as a solution to fill the gap between the low-cost, widely available computational resources and lack of user-friendly bioinformatics software. Laboratories lacking NGS data analysis expertise can easily perform analysis of data generated from in-house sequencing instruments or external service providers using this platform. The miCloud is highly modular and is based on Docker virtual machine containers [13] with components (e.g., user interface, file manager,pre-configured data analysis pipelines) encapsulated in separate containers (**Fig 1a**). These virtual machine containers are automatically instantiated and interconnected upon installation of the miCloud, which is as simple as running a script that users can download from our code repository ([14], **Suppl.**). As a result, users have access to an on-premises cloud application that leverages their local computational infrastructure and can access the miCloud graphical user interface (**Fig 1b**) via a web browser on a personal computer. The interface provides built-in functionality for users to run and monitor a set of preconfigured bioinformatics analysis pipelines, in addition to import, export, and management of NGS sequencing data through a visual file explorer (**Fig 1c**). By default, there are three pipelines ready to execute with the miCloud upon installation, two for single and paired-end ChIP-Seq data, in addition to one more for paired-end RNA-Seq data (**Suppl.**). Each is a multi-step pipeline processing the input data through a set of bioinformatics algorithms, which provide beginning to end data analysis and generate differential gene expression charts, tables of p-values for chromatin peaks found in the genome, and lists of SNP polymorphisms. The pipelines have been implemented following standard published protocols for RNA-seq and CHIP-seq [15], [16]. Additionally, we have integrated the Visual Omics Explorer (VOE, [17]) with the miCloud, to provide users with access to rich, interactive visualizations and publication-ready graphics from the pipeline outputs.

**Figure 1:**
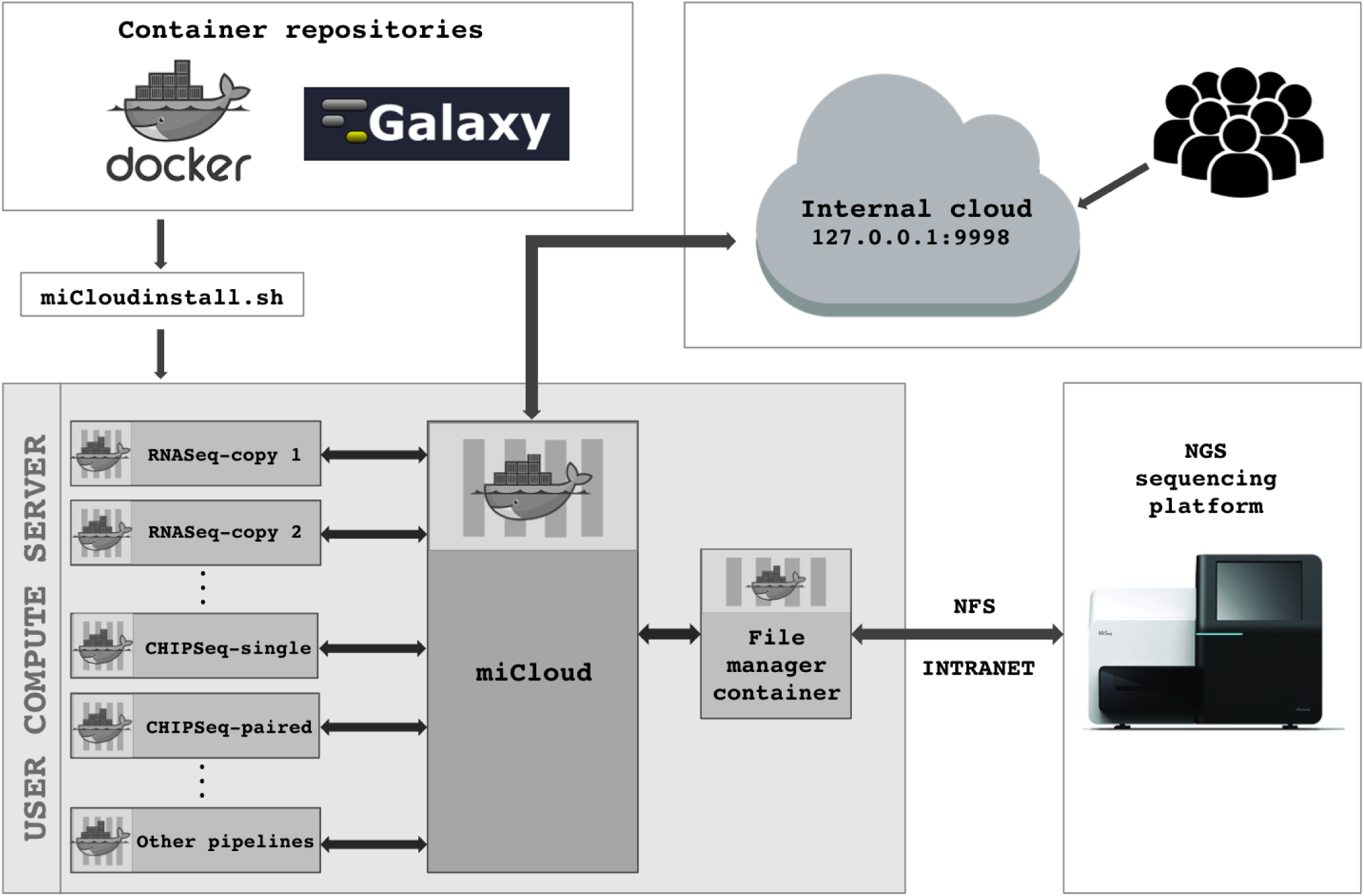
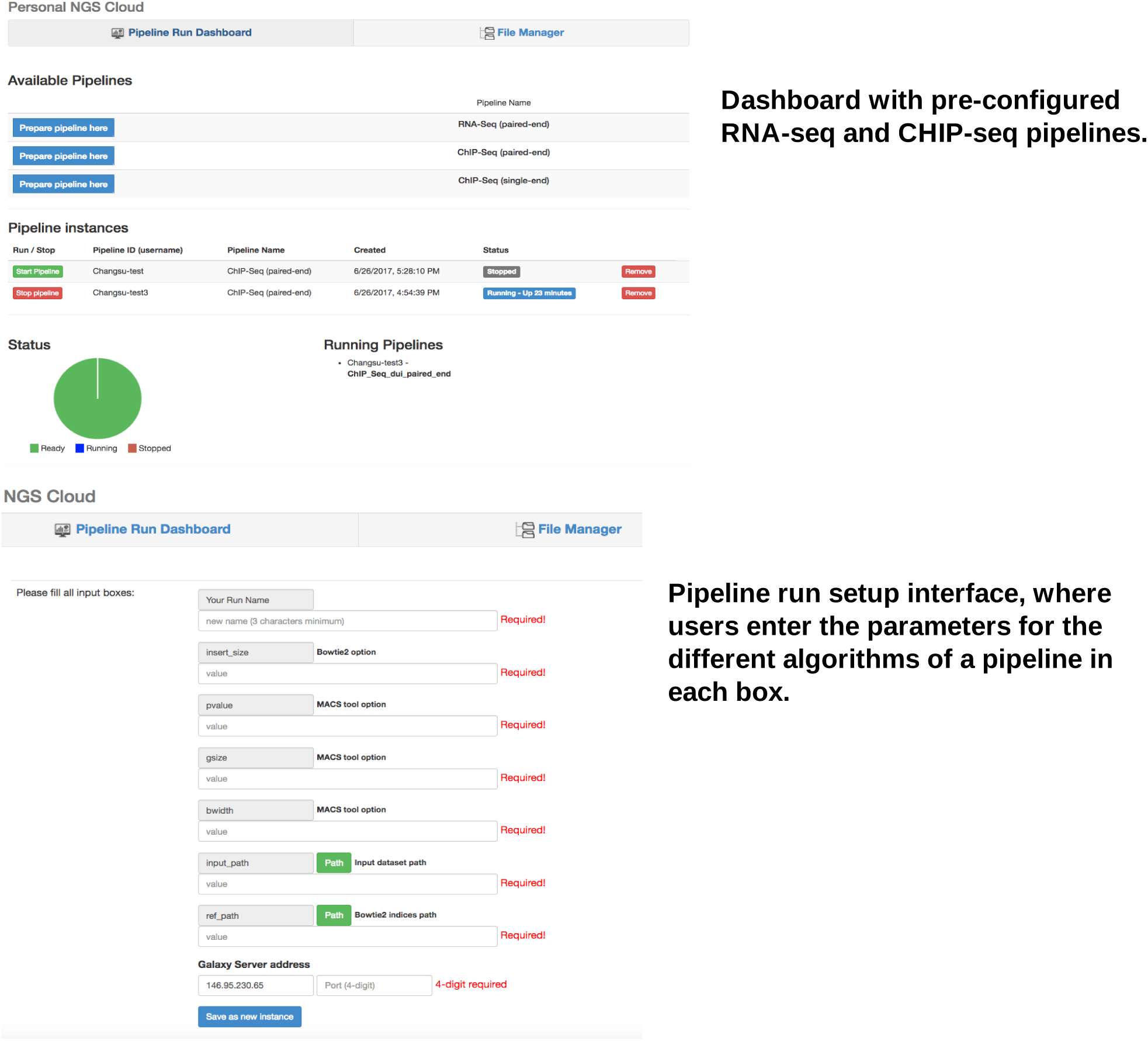
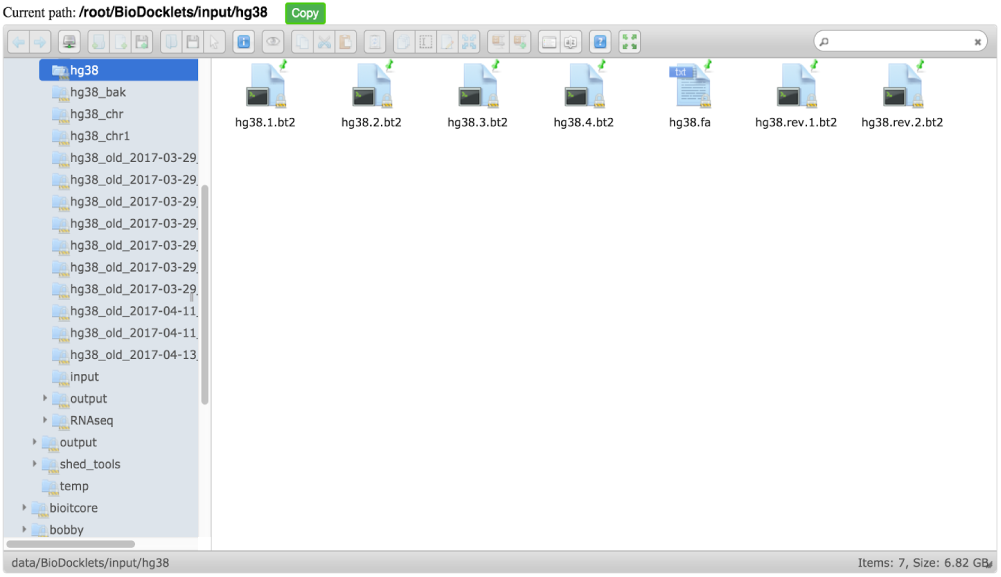
A. miCloud modular design with the different components in Docker containers. B. miCloud interface. C. miCloud file manager enabling users to browser local network file locations.

When the required computational capacity is available, users can start multiple replicas of each container through the miCloud interface, to run copies of the pipeline in parallel and process multiple datasets (**Suppl.**). The miCloud platform builds on technology we have previously developed through the Bio-Docklets project [18], and for the pipeline implementations, we have used the Galaxy workflow engine which runs inside the Docker containers. The BioBlend software library is also integrated with the miCloud, enabling seamless connection to the Galaxy Application Programming Interface (API). Furthermore, access to the API is used internally by the miCloud for controlling the execution of the pipelines within the Docker containers (**Fig 1c**), in addition to managing data inputs and outputs from the containers to the local filesystem. With these technologies as the foundation, we provide a fully extensible platform enabling developers to implement new Docker containers with bioinformatics pipelines and make them easily accessible for users through the miCloud interface. Developers can also leverage our platform to provide access to NGS data analysis containers publicly available from the Galaxy community [19] or Docker Store [20].

While Illumina sequencing instruments include a complete Windows desktop computer onboard that is used for the instrument control software, the operating system and data storage capacity of the onboard computer is not suitable for running the data-intensive pipelines such as RNA-seq and CHIP-seq. However, the operating system’s desktop can be accessed through the touch-screen monitor on the instrument [21], where users can simply create a Windows Network File System (NFS, [22]) folder with password protection, to share the sequencing read data (**Suppl.**) on the local network. The miCloud file manager was implemented so that it includes build-in NFS connectivity, allowing users to easily connect to the Illumina instrument and transfer the data over the local network on the computer or server where the miCloud containers are running.

The miCloud provides a bioinformatics platform for NGS data analysis that can be deployed without any technical expertise, enabling laboratories with an Illumina desktop genome sequencer, to seamlessly integrate the instrument with a fully-featured, on-site data analysis cloud. Bioinformatics developers can leverage the plethora of publicly available Docker containers with preconfigured NGS data analysis pipelines, to deploy complete data analysis solutions through miCloud. With this approach, non-bioinformatics experts have easy access to a fully-featured solution for execution of complex bioinformatics pipelines, where the underlying software complexity is fully abstracted through an intuitive interface.

